# A simplified transposon mutagenesis method to perform phenotypic forward genetic screens in cultured cells

**DOI:** 10.1101/598474

**Authors:** Charlotte R. Feddersen, Lexy S. Wadsworth, Eliot Y. Zhu, Hayley R. Vaughn, Andrew P. Voigt, Jesse D. Riordan, Adam J. Dupuy

## Abstract

The introduction of genome-wide shRNA and CRISPR libraries has facilitated cell-based screens to identify loss-of-function mutations associated with a phenotype of interest. Approaches to perform analogous gain-of-function screens are less common, although some reports have utilized arrayed viral expression libraries or the CRISPR activation system. However, a variety of technical and logistical challenges make these approaches difficult for many labs to execute. In addition, genome-wide shRNA or CRISPR libraries typically contain of hundreds of thousands of individual engineered elements, and the associated complexity creates issues with replication and reproducibility for these methods. Here we describe a simple, reproducible approach using the Sleeping Beauty transposon system to perform phenotypic cell-based genetic screens. This approach employs only three plasmids to perform unbiased, whole-genome transposon mutagenesis. We also describe a ligation-mediated PCR method that can be used in conjunction with the included software tools to map raw sequence data, identify candidate genes associated with phenotypes of interest, and predict the impact of recurrent transposon insertions on candidate gene function. Finally, we demonstrate the high reproducibility of our approach by having three individuals perform independent replicates of a mutagenesis screen to identify drivers of vemurafenib resistance in cultured melanoma cells. Collectively, our work establishes a facile, adaptable method that can be performed by *labs of any size* to perform robust, genome-wide screens to identify genes that influence phenotypes of interest.

## INTRODUCTION

Forward genetic screens, in which a phenotype of interest is selected from a population of mutagenized individuals, have long been viewed as a powerful tool to uncover novel components of biological systems. A variety of approaches have been used in model organisms such as yeast (Forsburg 2001), *Caenorhabditis elegans* (Jorgensen and Mango 2002), and fruit flies (St Johnston 2002). However, forward genetic screens have been more challenging to perform in mammalian organisms, in part due to the size and complexity of mammalian genomes. Chemical mutagenesis screens have been generally useful for obtaining interesting mutant phenotypes in mice, but the identification of the causative genetic alterations is laborious, even with the advent of genome sequencing.

The development of genome-wide shRNA and CRISPR libraries has facilitated cell-based screens to identify loss-of-function mutations associated with specific phenotypes. Hundreds of studies have been reported using either RNAi or CRISPR screens to identify genes associated with a wide variety of phenotypes (Schmidt et al. 2013; Rauscher et al. 2017), including extensive work to understand the vulnerabilities of cancer cell lines (Ling et al. 2018). Fewer options exist to perform gain-of-function (*e.g.* over-expression) genome-wide screens in cell-based assays. The typical approach utilizes arrayed lentiviral libraries consisting of hundreds to many thousands constructs, each expressing a single open-reading frame (ORF).

Concern remains regarding the consistency of such methods, given the substantial complexity involved with employing genome-wide libraries. Many screening libraries contain over 100,000 individual lentiviral constructs, which are typically synthesized and cloned into expression vectors in a pooled format. Inherent differences in the efficiency of vector propagation and packaging during these steps creates pools that lack homogeneity in terms of the quantity of each individual reagent. Production of arrayed libraries also requires substantial quality controls and automated liquid handling automation capabilities that most research facilities lack. Because of these issues, such genome-wide screens must be carefully designed and executed, including the use of complex statistical models to interpret and remove the substantial number of false positive hits. Ultimately, the complexity and expense associated with existing genome-wide screening approaches limits the ability of independent research groups to conduct novel screens or replicate previously-reported results.

Compared to complex genome-wide screening methods that individually target elements at the genome scale, insertional mutagenesis screens are generally much simpler. Retroviral insertional mutagenesis has been used to select for phenotypes and mutations of interest in cultured cells (Kandel et al. 2005; Gragerov et al. 2007; Kizilors et al. 2017). However, retroviral vectors typically exhibit significant insertion bias, and proviral integration can have complex effects on gene expression, thus limiting the utility of viral insertional mutagenesis. By contrast, transposon systems, such as Sleeping Beauty (SB) and piggyBac, have become more commonly used for insertional mutagenesis due to their flexible design and reduced integration site bias. While transposon mutagenesis has been used to perform phenotypic selection in cells *ex vivo* (Chen et al. 2016; Guo et al. 2016; Kodama et al. 2016), it has more frequently been employed in engineered mouse models of cancer (O’Donnell 2018), likely due to the relative inefficiency of mutagenesis when both transposon and transposase vectors must be introduced independently to cells in culture.

Here we describe a novel method to perform simple, phenotype-driven genome-wide genetic screens in cultured cells using a hyperactive version of the Sleeping Beauty transposon system. Unlike other genome-wide screening methods, ours consists of only three plasmids that are used to carry out mutagenesis in cultured cells. Following phenotypic selection of mutagenized cells, a simple ligation-mediated PCR method (Supplemental Methods) prepares libraries that can be multiplexed and directly sequenced on the Illumina platform. We developed a series of software tools to take raw sequence data as input and perform trimming, mapping, and filtering of transposon integration sites. We also report an updated gene-centric (biased for analysis of annotated genes) common insertion site (gCIS) analysis tool modified from a previously described method to analyze transposon integration data from SB models of cancer (Brett et al. 2011) that predicts the functional impact of transposon insertions on the adjacent genes. Collectively, our methods and analytical tools provide labs of virtually any size with the ability to independently perform genome-wide phenotypic screens in cultured cells.

## RESULTS

### Transposon mutagenesis in cultured cells

We considered several factors when setting out to develop a cell-based forward mutagenesis screening approach. Ideally, a plasmid-based system provides the greatest flexibility to mutagenize any cell line that can be transfected with moderate to high efficiency. While our initial efforts with the SB11 transposase enzyme (Geurts et al. 2003) did not provide sufficient mutagenesis efficiency for screens (not shown), a hyperactive version of the SB transposase with ∼100-fold increased activity, SB100X (Mates et al. 2009), was sufficient for screening purposes. We generated a piggyBac vector to create a simple, viral-free method to establish stable expression of the SB transposase in mammalian cells by simply co-transfecting the SB100X vector with a second vector encoding a hyperactive version of the piggyBac transposase (hyPBase)(Yusa et al. 2011)(**Fig.1**). Once cells stably expressing SB100X are generated, transposon mutagenesis is performed by simply transfecting a mutagenic transposon (*e.g.* pT2-Onc, pT2-Onc2, pT2-Onc3) (Collier et al. 2005; Dupuy et al. 2005; Dupuy et al. 2009). The SB100X enzyme then mobilizes mutagenic transposons from the transfected plasmid into the cellular genome. Once integrated, the transposons can impact gene expression through a few known mechanisms (Dupuy 2010).

**Figure 1.**
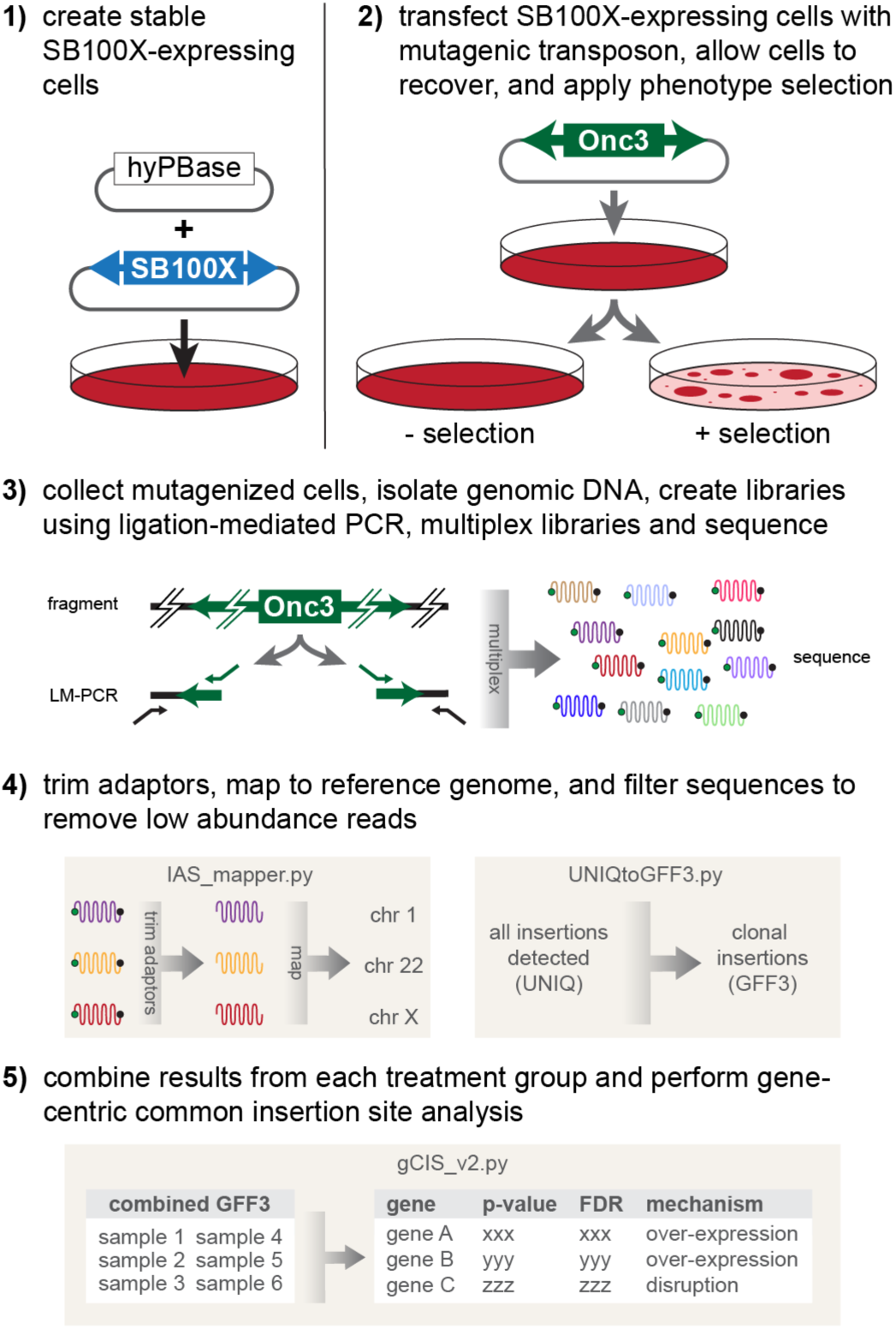
Overview of cell-based Sleeping Beauty (SB) mutagenesis screens. (**1**) Cells are engineered to express the SB100X transposase. This is done using a piggyBac transposon expression system using the hyPBase transposase to stably deliver the SB100X transgene. (**2**) Once SB100X-expressing cells are selected, cells are transfected with the T2-Onc3 plasmid, or similar mutagenic SB transposon vector, and cells with the desired phenotype are selected from among the population of mutagenized cells. (**3**) Genomic DNA is collected from each population of selected and control cells, and each DNA sample is subjected to a ligation-mediated PCR approach to isolate the transposon-genome junctions. Individual LM-PCR libraries are then multiplexed and sequenced on the Illumina platform. (**4**) Raw sequences are trimmed, mapped to the reference genome, and filtered based on user-defined criteria. (**5**) Finally, filtered sequences are combined and analyzed to identify candidate genes enriched for transposon insertions in selected but not control cells.

To characterize our forward mutagenesis screening system, we performed a screen to identify novel drivers of resistance to the BRAF inhibitor vemurafenib in cultured melanoma cells. We selected the A375 melanoma cell based on its recurrent usage in shRNA/sgRNA screens to identify drivers of vemurafenib resistance (Whittaker et al. 2013; Shalem et al. 2014; Joung et al. 2017). A population of A375 cells stably expressing SB100X transposase (A375-SB100X) was produced and subsequently transfected with either the mutagenic pT2-Onc3 transposon vector (Dupuy et al. 2009) or a control EGFP expression plasmid. Cells were grown for 48 hours to facilitate integration of mutagenic transposons, independent plates of mutagenized or control cells were pooled, and 1 × 105 cells were seeded onto 6-well plates. Twelve to twenty-four hours post-plating, cells were placed under drug selection in 3, 5, or 10 µM vemurafenib or vehicle control (dimethylsulfoxide). Media was changed twice weekly, and wells were monitored for the emergence of resistant colonies. Both mutagenized and control plates were harvested and stained when resistant colonies were plainly observable in control wells.

Pilot experiments indicated that 5 µM vemurafenib was the optimal drug concentration to select for resistant SB-mutagenized A375 cells, based on several observations. At 3 µM vemurafenib, resistant colonies emerged at roughly the same time in treated and control wells, indicating an insufficient stringency to distinguish between spontaneous resistance and resistance driven by transposon mutagenesis. While 10 µM vemurafenib greatly reduced the frequency of spontaneous resistance in the control wells, colonies that eventually emerged in mutagenized cells were fewer and smaller than those that grew at lower drug concentrations. In contrast, at 5 µM vemurafenib, mutagenized A375 cells produced large colonies in only 10-14 days, while control cells did not produce large resistant clones in the same timeframe (**Fig.2**). Based on these results, we selected a 5 µM vemurafenib dose to perform our mutagenesis screen for drivers of vemurafenib resistance.

**Figure 2.**
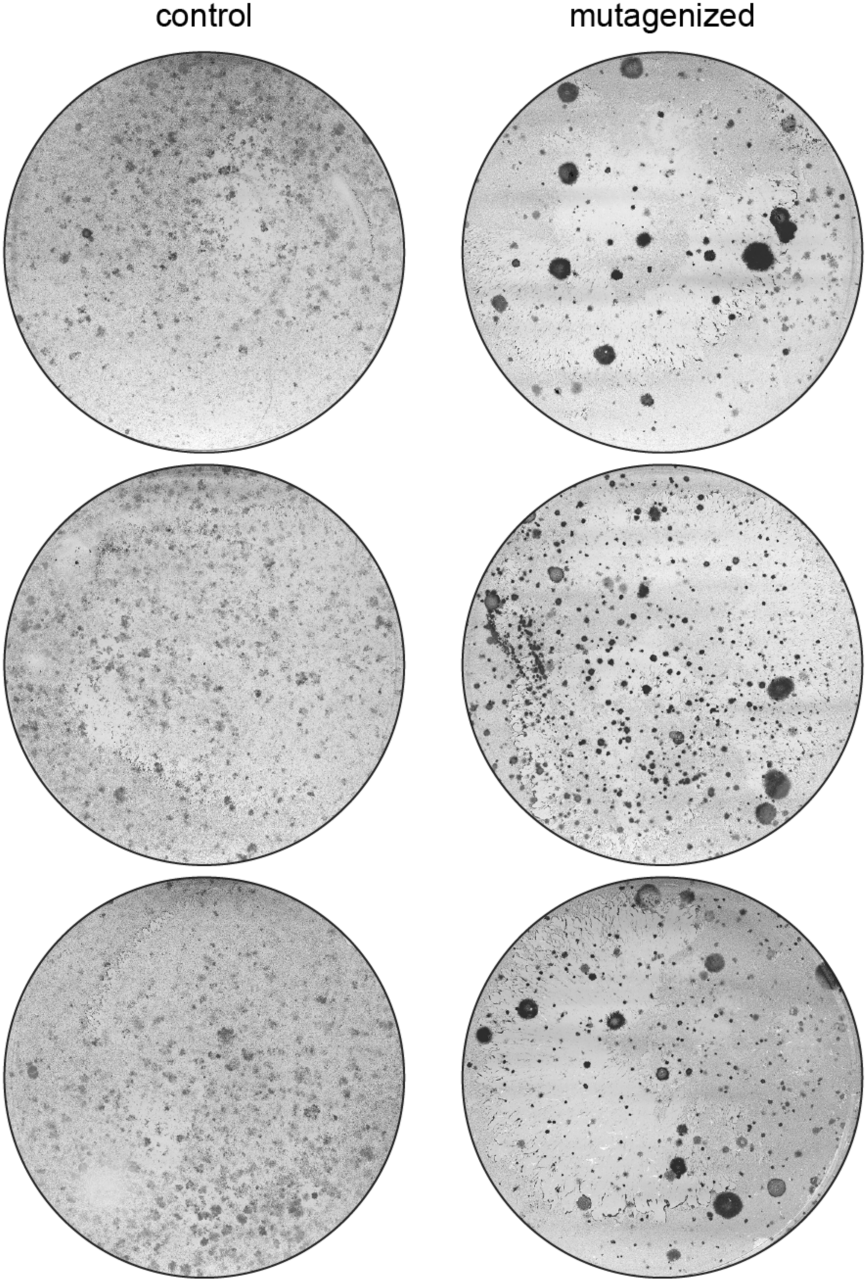
Transposon mutagenesis drives vemurafenib resistance in A375 cells. Cells were engineered as described in Figure 1. Twenty-four hours after mutagenesis was initiated, 10 cm plates were seeded with mutagenized or control cells. After allowing cells to attach, 5 µM vemurafenib was added to the culture medium. Culture media and drug were replaced twice weekly until colonies emerged on the plates containing mutagenized cells. Colonies were stained with Coomassie Blue. Representative plates are shown.

A375-SB100X cells were transfected with the pT2-Onc3 transposon plasmid to initialize mutagenesis. Twenty-four hours post-transfection, 10-centimeter plates were seeded with 1 x 10^6^ cells and allowed to recover for 12-24 hours. After recovery, each plate was treated with either 5 µM vemurafenib or dimethylsulfoxide (*i.e.* vehicle control). While vehicle-treated cells expanded rapidly and reached confluency ∼3 days after plating, drug-resistant colonies emerged within 10-14 days after the addition of drug, as previously observed in the pilot study. Importantly, this experiment was independently conducted by three different members of the lab with similar results (see Methods). In all cases, vemurafenib resistant colonies emerged in mutagenized cells in 10-14 days while control cells did not develop spontaneous resistance in the same timeframe.

### Identification of transposon insertion sites

Next, we sought to determine if recurrent transposon-induced mutations could be identified in resistant colonies. As picking individual colonies would significantly limit throughput, we instead harvested independent plates of cells as pooled populations, as is typically done with genome-wide shRNA and CRISPR screens. We collected pooled cell populations from each independent 10 cm plate and extracted genomic DNA. Fragments containing the transposon/genome junctions were amplified via ligation-mediated PCR (LM-PCR) and sequenced using the Illumina HiSeq 4000 platform (Supplemental Methods). A single FASTQ file was generated for each population of vemurafenib-(n=69) and vehicle-treated (n=15) cell population. We generated an informatics pipeline (IAS_mapper) to processes each FASTQ file, trimming the residual transposon and adaptor sequences and mapping trimmed reads to the human reference genome (GRCh38). Although we designed the IAS_mapper pipeline to utilize adaptor trimming (cutadapt)(Martin 2011) and mapping (HISAT2)(Kim et al. 2015) methods that can be run on a standard workstation, the software can be modified to run on a parallel processing environment to decrease processing time.

### SB100X-mediated transposition results in numerous integration events per cell

We sought to determine the efficiency of mutagenesis at the cellular level to get an estimate of the number of insertion events that can be achieved with our approach. To do this, we isolated and expanded eight individual vemurafenib-resistant colonies derived from three independent plates. Following expansion, we verified the drug-resistance of a subset of colonies in a 10-day growth assay (**Supplemental Fig.1**). We then identified transposon integration sites as described above. Interestingly, the number of insertions within each colony ranged from 10 to 135 (**Fig.3**). Colonies harboring large numbers of insertions are likely to be mixed populations due to collection and isolation process. However, it is also possible that single cells harbor >100 transposon insertion events, similar to Sleeping Beauty-induced leukemias we have previously described (Berquam-Vrieze et al. 2011). Regardless, these results indicate that transposon mobilization from transfected plasmids is highly efficient, resulting in numerous insertional mutation events per cell.

**Figure 3.**
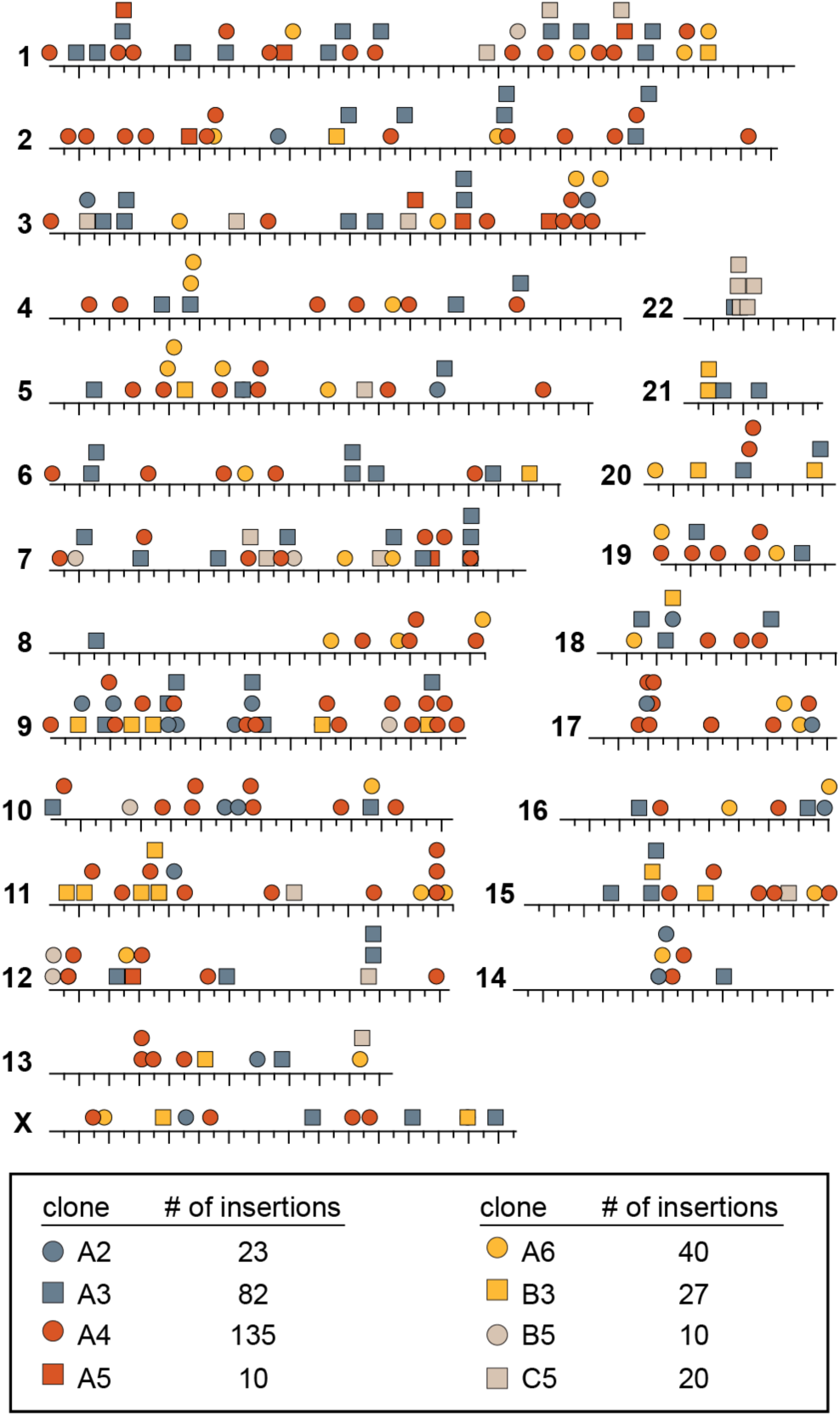
Comparison of transposon insertion sites in a series of vemurafenib-resistant A375 colonies. Eight distinct vemurafenib-resistant colonies were isolated from three independent populations of SB-mutagenized A375 cells. Each colony was expanded and transposon insertion sites were identified (see Figure 1). Between 10 and 135 insertion events were present in each expanded colony. In general, transposon insertion sites were distributed across the genome, suggesting that transposon mutagenesis is capable of performing genome-wide screens.

### Gene-centric common insertion site analysis identifies recurrently mutated genes in selected cells

The identification of candidate genes in SB screens essentially relies on the identification of genes that suffer transposon insertion events at a rate that exceeds the expected background rate of mutation, based on the experimental parameters. While this concept is straightforward, most integrating vector systems display insertion site bias, often favoring regions near transcription start sites. Therefore, integration site bias can produce a high rate of false-positive gene candidates. A variety of prior studies have shown that the SB transposase exhibits little integration site bias relative to other integrating vectors, requiring only a TA dinucleotide for insertion (Vigdal et al. 2002; Yant et al. 2005; Liang et al. 2009; Ammar et al. 2012; Woodard et al. 2012; Rogers et al. 2013; Riordan et al. 2014). This feature of the SB system is beneficial because it 1) allows SB mutagenesis to access virtually the entire genome and 2) simplifies the analysis of SB integration data by eliminating the need to produce complex statistical models that account for bias.

We have previously described a method to identify genes that harbor recurrent SB transposon insertion events in association with a phenotype of interest (Brett et al. 2011). This method, referred to as gene-centric common insertion site (gCIS) analysis, performs a simple chi-squared test to compare the number of observed insertion events in a given gene to the number of expected insertion events based the number of TA sites within the gene and the total number of insertion events identified in the data set. The initial gCIS method was designed to analyze insertion data derived from SB-induced tumors in engineered mouse models. As a consequence, the pipeline assumes each analyzed sample is a single, clonally-derived cell population (*i.e.* a tumor), not a pooled population of unrelated clones.

We developed a new gCIS method (gCIS2) to analyze transposon insertion data from cell-based phenotypic screens, such as our vemurafenib resistance screen performed in A375 melanoma cells (for detailed description see Supplemental Methods). Given that we used a pooled sample preparation to facilitate higher throughput for the screen, we were unable to determine which insertion sites co-occur within the same cell clone. Therefore, we evaluated each unique insertion site as an independent event. This approach ignores potential interactions between insertions that co-occur in the same cell and instead assumes that single insertion events can provide the desired phenotype, allowing evaluation of cell pools containing an unknown number of clonal populations.

We next analyzed insertion site data from the vemurafenib-and vehicle-treated A375 cell populations using the gCIS2 tool (**Supplemental Table 1**). The pipeline identified 11,094 unique transposon insertion events in 7,598 RefSeq genes, including the transcription unit and promoter region - defined as 40 kilobases upstream of the start site. Overall, nine genes were significantly enriched for transposon insertions (false discovery rate ≤ 1 × 10^-5^), were unique to the vemurafenib-treated population (see Methods) and showed a transposon distribution pattern and orientation trend consistent with either over-expression or disruption. For example, the pipeline identified 162 independent transposon insertions within the 5’ region of the *VAV1* locus. Nearly all of the transposons (161/162) inserted in the promoter region or first intron of the locus, and the vast majority (88%) of transposons inserted in the same transcriptional orientation as the *VAV1* gene, suggesting that over-expression of VAV1 is associated with vemurafenib resistance in A375 cells.

### Functional prediction of transposon effects on candidate genes

Prior insertional mutagenesis screen analysis pipelines, including our original gCIS method, identify candidate genes by searching for clusters of transposon insertion events within the genome but do not attempt to predict the impact of transposon insertion on the affected genes. Mutagenic SB transposons were developed to modify expression of endogenous genes in specific ways depending on their relative position and orientation, as previously described (Dupuy 2010). This feature, combined with our experience analyzing insertion site data from 15 SB-induced models of cancer (Berquam-Vrieze et al. 2011; Mann et al. 2012; Wu et al. 2012; Keng et al. 2013; Quintana et al. 2013; Rogers et al. 2013; Zanesi et al. 2013; Bard-Chapeau et al. 2014; Mann et al. 2015; Takeda et al. 2015; Montero-Conde et al. 2017; Morris et al. 2017; Suarez-Cabrera et al. 2017; Tschida et al. 2017; Riordan et al. 2018), led us to develop an algorithm to predict the functional impact that recurrent transposon insertion has on a given gene. This approach evaluates clustered transposon insertions within each candidate locus to determine if there is a bias for insertion in the same orientation as the gene, indicative of an over-expression mechanism, or if the transposons are randomly orientated, suggesting a gene disruption mechanism (see Supplemental Methods). Of the nine genes identified by gCIS2, eight were predicted to drive resistance through over-expression and one was predicted to drive resistance via gene disruption (**Supplemental Table 1**). Importantly, we have recently described experiments that functionally validated the role of four of these candidates (VAV1, MCF2, BRAF, RAF1) in driving vemurafenib resistance in human melanoma cell lines (Feddersen et al. 2019).

We next evaluated the eight genes predicted to drive vemurafenib resistance via over-expression, as determined by gCIS2. Transposon orientation bias was evident in each of these genes, and in the majority of cases the cluster of transposon insertions was centered in the 5’ region. For two genes, *BRAF* and *RAF1*, transposons were clustered within the gene body well downstream of the promoter, a pattern predicted to drive over-expression of an mRNA encoding an N-terminally truncated protein (**Supplemental Fig.2**). We modified the gCIS2 method to facilitate the identification of rare genes in which a similar protein truncation mechanism may be occurring, evaluating the distribution of transposon insertion sites across each gene to calculate the skewness and kurtosis (**Fig.4A**). Skewness provides an indication of clustering near the 5’ (positive skewness) or 3’ (negative skewness) region of the gene. Kurtosis provides an indication of how tight (positive kurtosis) or loose (negative kurtosis) the distribution of transposon insertions is within the gene (**Fig.4A**).

**Figure 4.**
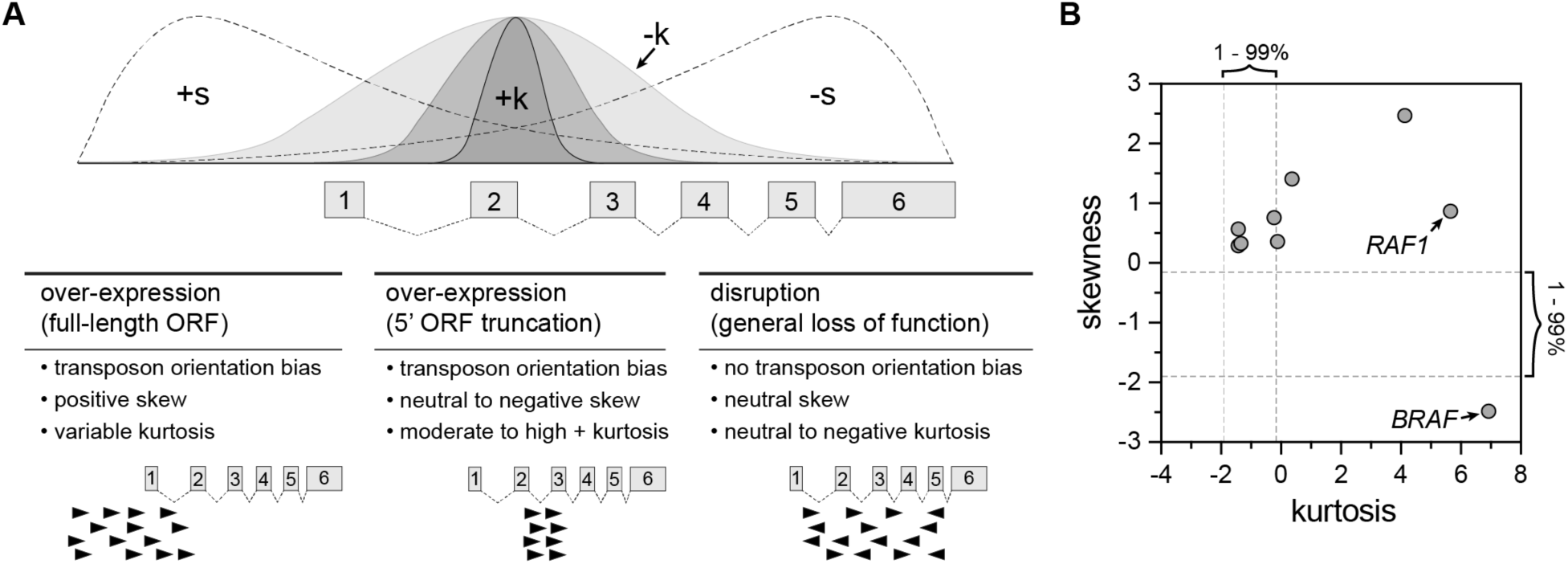
Predicting the functional impact of transposon insertion on individual candidate genes. The gCIS2 pipeline employs an algorithm for each gene that has ≥ 5 independent insertions to predict the impact of transposon insertion on the gene (i.e. over-expression or gene disruption). (**A**) The gCIS2 pipeline also determines the skewness and kurtosis values for the distribution of transposon insertions across the gene. These values can be evaluated by the user to further distinguish between different mechanisms of over-expression (*i.e.* full-length vs. 5’truncation). (**B**) Actual skewness and kurtosis values are shown for the candidate drivers of vemurafenib resistance identified in A375 cells. [dashed lines indicate the 1-99% intervals for skewness and kurtosis obtained from the analysis of 20 independent simulated data sets (see Methods).]

### Evaluation of gCIS2 performance using simulated data sets

We have previously compared SB transposon integration data to matched sets of randomly selected sets of TA dinucleotide sites to detect non-random patterns in the experimental data set (Riordan et al. 2014). Using this approach, we generated twenty independent, simulated data sets by replacing each transposon insertion event with a randomly selected TA dinucleotide site from the human reference genome (GRCh38). Each simulated data set was then analyzed by gCIS2.

As described above, gCIS2 identified nine candidate resistance genes in the experimental data set that met the criteria defined by the pipeline (*i.e.* FDR ≤ 1 × 10^-5^ with conserved function prediction). By contrast, gCIS2 did not identify genes that met the same criteria in the twenty simulated random data sets. This result is not surprising given that the gCIS2 method assumes a random distribution of transposon insertion sites. Next, we compared skewness and kurtosis values between the experimental and simulated data sets.

Next, we compared skewness and kurtosis values between the experimental and simulated data sets. We determined the mean skewness and kurtosis values for the 1^st^ and 99^th^ percentiles among the twenty simulated data sets (**Supplemental Fig.3**). Generally, the experimental data set had limits similar to the simulated data sets for both skewness and kurtosis but had significantly greater values, particularly for kurtosis. This result demonstrates that the experimental data set contains more tightly clustered transposons within some genes, indicative of strong selective pressure for integration within a relatively small genomic region within each locus.

The gCIS2 pipeline calculates both skewness and kurtosis for each gene analyzed, but these values are not used to make predictions about the functional impact of transposon insertion on a given gene. Instead, they can be used to evaluate the extent to which transposons show a conserved functional mechanism such as over-expression of a full length open-reading frame (ORF), over-expression of a truncated ORF, or gene disruption (**Fig.4A**). For example, the skewness and kurtosis values calculated for each of the nine candidate genes identified by gCIS2 in the experimental data set shows particularly high kurtosis values for both *RAF1* and *BRAF* (**Fig.4B**). This feature could be useful in identifying cases in which transposon-induced gene truncation is associated with the selected phenotype, vemurafenib resistance in this case.

### Independent replicates of vemurafenib-resistance screen yield nearly identical results

A major criticism of genome-wide screens using shRNA and CRISPR is the lack of reproducibility between independent experiments. For example, three independent genome-wide loss-of-function screens to identify drivers of vemurafenib resistance in A375 cells have been reported (Whittaker et al. 2013; Shalem et al. 2014; le Sage et al. 2017). Of the 34 candidate genes reported in these publications, only *NF1* was identified by all three. As previously mentioned, three different individuals in this study performed our vemurafenib resistance SB mutagenesis screen in A375 cells using the same procedure. In part, this strategy was used to determine the reproducibility of our transposon screening method. Analysis of the entire data set identified nine candidate genes (**Supplemental Table 1**). Encouragingly, 6 of these 9 genes were identified as significant in all three replicates of the screen, despite significant variations in the sample size (**Fig.5**). The remaining three genes were identified as significant in two of three screens. This finding provides evidence that the transposon mutagenesis method described here provides a high degree of reproducibility.

**Figure 5.**
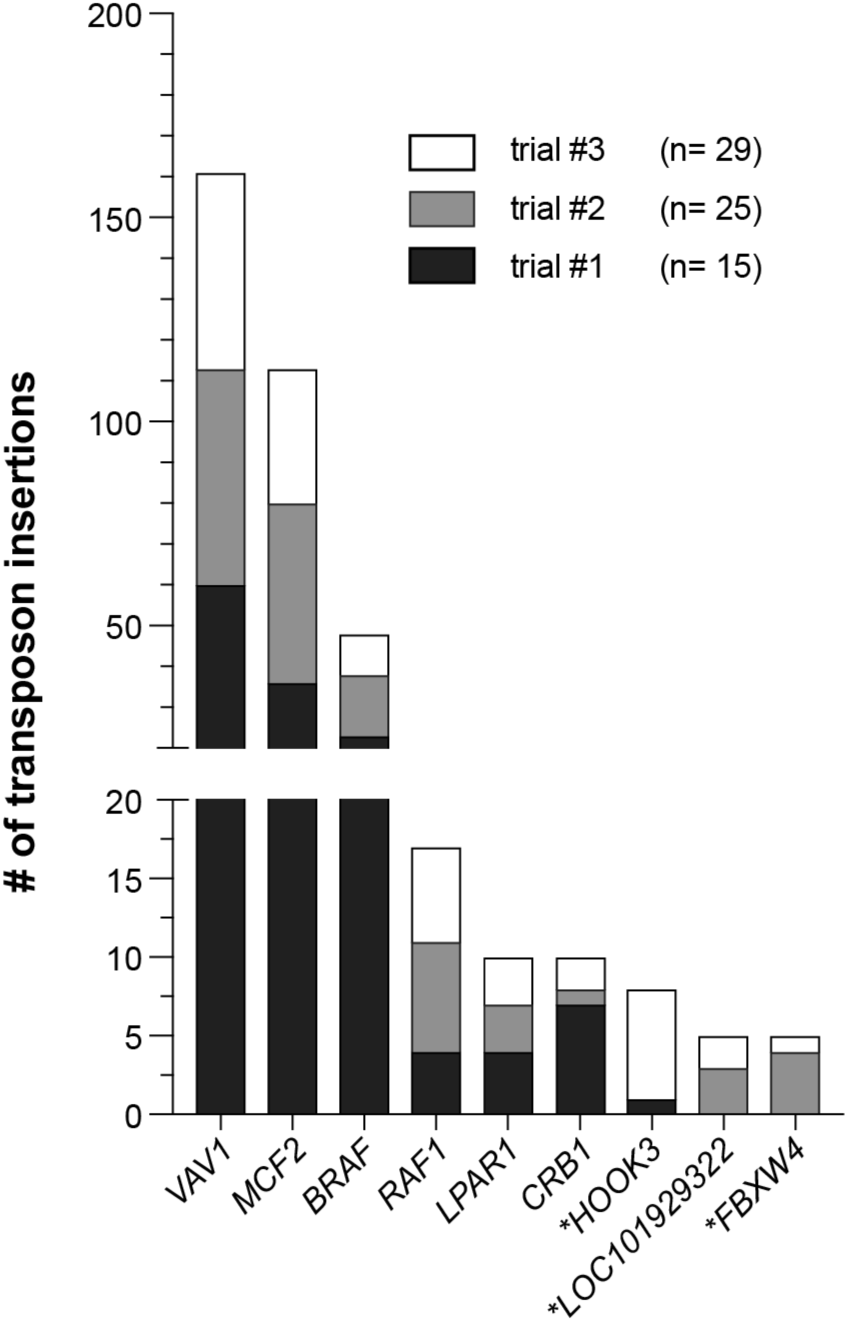
High degree of replication in cell-based SB mutagenesis screens. The vemurafenib resistance screen was replicated by three different individuals spanning a period of several months. The gCIS2 results were obtained for each independent replicate and compared to the results obtained by analyzing the combined data set. Of the nine vemurafenib resistance driver genes identified by analyzing the entire data set, six were identified by analyzing each replicate independently while the remaining three genes (indicated with asterisks) were identified in two of the three replicates.

## DISCUSSION

Here we describe the development of both the molecular and bioinformatic tools needed to perform Sleeping Beauty (SB) transposon-based phenotypic screens in cultured mammalian cells. Previously, the application of SB transposon mutagenesis was mostly restricted to genetically engineered mouse models of cancer. Our prior attempts at performing cell-based mutagenesis using earlier versions of the SB transposase failed to produce vemurafenib-resistant A375 cells at a frequency above the spontaneous rate of resistance (not shown), presumably due to insufficient transposition efficiency. However, we show here that the use of the hyperactive SB100X transposase mediates multiple transposon insertion events per cell when transposons are introduced as plasmids via standard transfection (**Fig.3**).

When designing this method, we considered the major technical and logistical limitations of current genome-wide genetic screening techniques (*e.g.* shRNA, CRISPR). Our goal was to develop a system that is simple, cost-effective, and reproducible. Importantly, we also wanted to provide the bioinformatic tools necessary to enable labs of virtually any size to perform mutagenesis screens and analyze the data using the methods we describe here.

Our first application of this mutagenesis approach was to derive human melanoma cells resistant to the BRAF inhibitor vemurafenib (**Fig.2**)(Feddersen et al. 2019). This screen identified nine candidate genes as drivers of vemurafenib resistance in A375 cells (**Supplemental Table 1**). A major advantage of our SB mutagenesis approach is its methodological simplicity, as demonstrated by the ability of three different individuals to perform the screen and obtain highly similar results (**Fig.5**). This outcome was achieved with the methods described here in detail using commercially-available reagents.

This flexible approach can be easily adapted to identify genes associated with any selectable phenotype. In general, we envision the use of our transposon-based screening method to work in the following manner. The process of generating mutagenized cells would stay relatively similar across experiments, with slight variations to achieve optimal transfection in the cell line being used. After mutagenesis is performed, the selection mechanism can be changed to fit the investigators’ biological question. A drug selection, like we describe here, is a simple application. Nevertheless, other selection mechanisms could be employed like cell sorting based on one or more parameters, such as expression of a reporter gene or surface marker, cell motility or invasion, or rescue experiments to restore cell viability following a particular insult. While the potential phenotypic screens that could be performed using this approach are seemingly endless, each screen will require optimization. There are a few areas to be considered when designing a new screen.

The selection method must be optimized to reduce the background signal in non-mutagenized cells (*i.e.* reduce the rate of spontaneous phenotype acquisition). For example, we initially conducted an experiment to determine the optimal concentration of vemurafenib that was high enough to slow the development of spontaneous drug resistance in A375 cells while still allowing for the emergence of resistant colonies in the mutagenized cell population (**Fig.2**). Choosing the appropriate selection stringency is extremely important and will vary by screen. Parameters to consider when choosing a selection dose are the rate of spontaneous phenotype acquisition and the hypothesized frequency of phenotype driver mutations. If the spontaneous rate of phenotype acquisition is high, increasing the stringency or lowering plating density can help prevent the emergence of spontaneous phenotype acquisition (*i.e.* non-transposon driven). Conversely, if the hypothesized frequency of mutation is low, lowering stringency may allow for weaker drivers to survive and proliferate. Regardless, the selection strategy for each screening method must be carefully considered.

Control cell populations must be included in each screen, and their production should be carefully considered. In addition to subjecting non-mutagenized controls to selection, which provides an assessment of the rate of spontaneous phenotype acquisition, a cell population that is mutagenized in the absence of selection is required. Ideally, these control cells should be derived from the same population of mutagenized cells that undergo phenotypic selection (see step 2 in **Fig.1**). To the greatest extent possible, cells in the control and experimental populations should be treated identically, aside from the phenotypic selection present in the experimental population. With this design, the control cells will facilitate the identification of false-positive gene hits (*i.e.* mutations that are enriched in the absence of the experimental selection). These false-positive gene hits can emerge from a variety of sources such as indirect selection for mutations that drive cell proliferation and/or viability, the production of PCR artifacts during the LM-PCR step prior to sequencing, and mapping artifacts produced during the sequence analysis step. Regardless of the source, analysis of the control cell population will identify genes that are not specifically associated with the phenotype.

Forward genetic screens allow for the association of specific genes with phenotypes of interest, and they have been indispensable in furthering our understanding of biology. While genome-wide approaches that employ RNAi or CRISPR have facilitated more rapid, cell-based screens, the complexity of performing and analyzing these experiments is beyond the capabilities of most labs. We provide here the reagents and tools necessary for virtually any lab to perform cell-based Sleeping Beauty screens.

## METHODS

### Mutagenesis of cultured A375 cells

Cells were stably transfected via Effectene (Qiagen) coupled with a piggyBac transposase integration system (Li et al., 2013) consisting of a piggyBac transposase-expressing plasmid and an Ef1α-SB100x transgene embedded within a piggyBac transposon. After puromycin selection, SB100X-positive cells were transfected with the pT2/Onc3 transposon plasmid (Dupuy et al., 2009), and 1 ×10^6^ mutagenized cells were plated on 10 cm plates 48 hours later. Cells were subsequently treated with vemurafenib or vehicle (DMSO, 0.2%) 24 hours after plating. Drug/vehicle was renewed every 3-4 days. Upon reaching confluency (approximately 3 days post-plating), vehicle plates were collected via trypsinization. This process was repeated independently at three different times by Hayley Vaughn (trial #1, n=15), Andrew Voigt (trial #2, n=25), and Eliot Zhu (trial #3, n=29).

For clonal isolation, a pipette tip was lightly placed in the center of a resistant colony and then dipped in media contained in a 24-well dish. Six colonies from three plates were collected in this manner for a total of 18 independent colonies. Two colonies failed to grow during the expansion phase and only eight were genetically profiled.

To identify common transposon insertion sites across plates of resistant cells or from expanded clonal populations, genomic DNA from each plate was collected via GenElute™ Mammalian Genomic DNA Miniprep Kit (Sigma). DNA fragments containing transposon/genome junctions were amplified via ligation mediated PCR and sequenced using the Illumina Hi-Seq 4000 platform.

### Colony staining

SB100X-A375 cells (1×10^5^) transfected with either pEGFP or pT2/Onc3 were stained with Coomassie Brilliant Blue after ethanol fixation after 25 days of drug treatment. Colonies were counted with the GelCount™ Colony Counter (Oxford Optronix).

### Cell culture

All cells were grown in DMEM (Gibco) supplemented with penicillin/streptomycin (Gibco) and 10% FBS (Atlanta Biologicals).

### Generation of simulated data sets

Independent data sets were generated by randomly selecting TA sites on each chromosome. The size of each random TA set was matched to the observed number of insertion events on each chromosome in the experimental data set.

### Access to analysis tools

All tools, including accompanying documentation, described in this manuscript can be acquired through GitHub (https://github.com/addupuy/IAS.git).

## DATA ACCESS

(including public database accession numbers for all newly generated data and/or reviewer links to deposited data when accessions are not yet public. Previously published accessions should be included in the Methods section where appropriate)

