## Supplemental Figures 1-3 for "A simplified transposon mutagenesis method to perform phenotypic forward genetic screens in cultured cells"

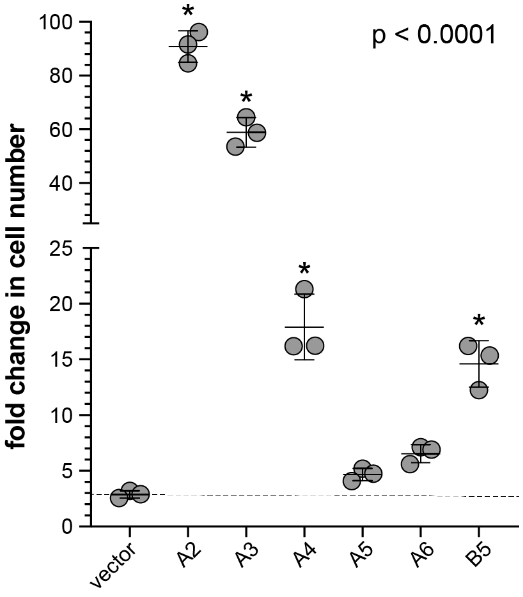


**Supplemental Figure 1.** Evaluation of expanded colonies for vemurafenib resistance. Six of the eight colonies were expanded and grown in a 96-well format in the presence of 5 µM vemurafenib. Cell number was indirectly measured using a CellTiter-Blue assay at day 0 and day 10. The fold change in signal is shown for each colony tested. An ANOVA test with a post-hoc analysis was performed to compare each colony to A375 cells carrying an empty expression vector. Each colony that is significantly different from the vector reference is indicated with an asterisk.

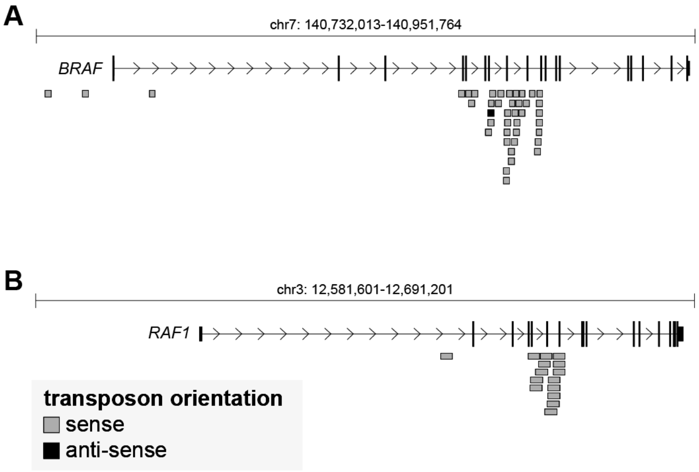


**Supplemental Figure 2.** Position of transposon insertions that generate N-terminal truncations of either *BRAF* (**A**) or *RAF1* (**B**) in vemurafenib-resistant A375 melanoma cells.

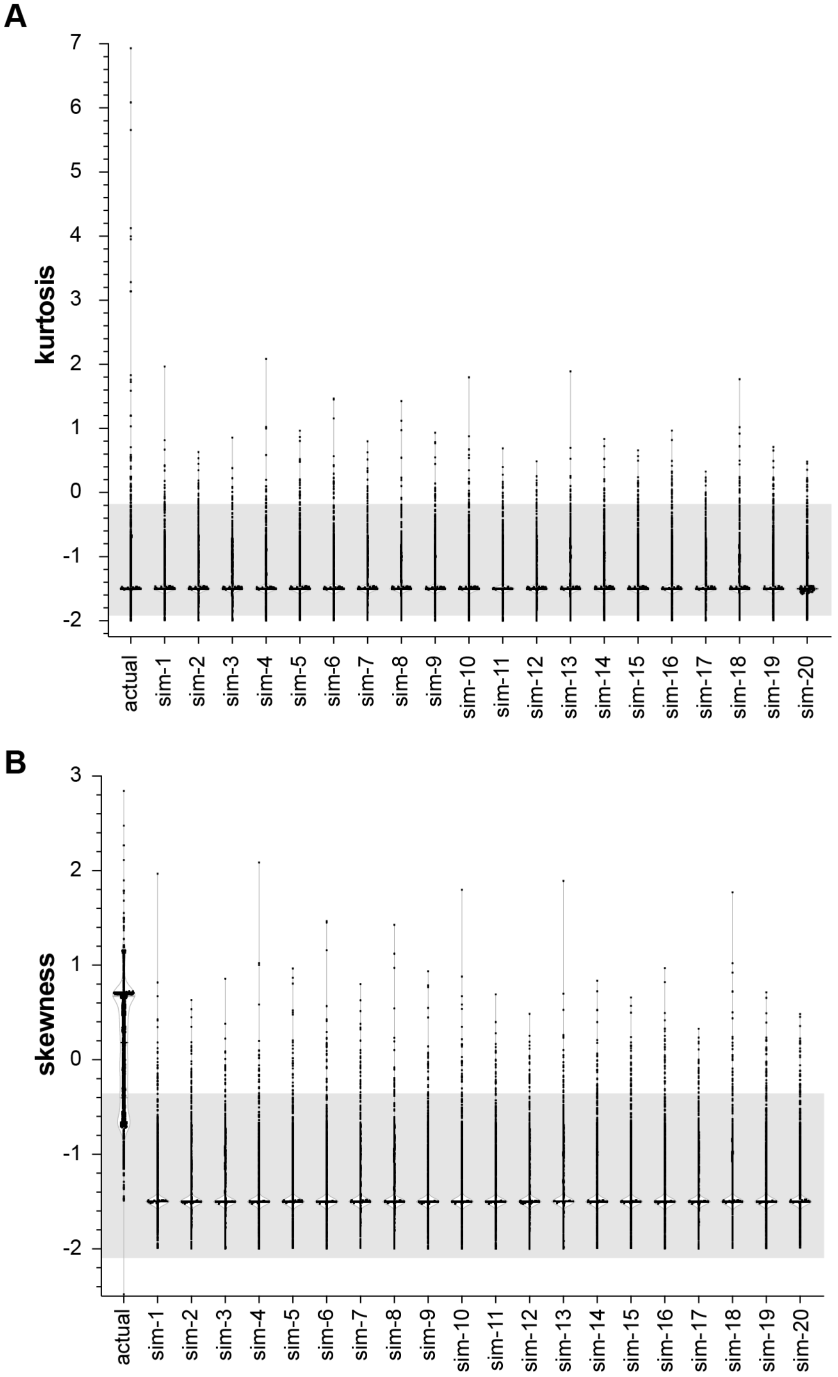


**Supplemental Figure 3.** Comparison of transposon cluster distribution measures of kurtosis (**A**) and skewness (**B**) between data derived from vemurafenib-resistant A375 cells and twenty independent simulated data sets. The total number and chromosomal distribution of the transposon insertions is matched for each simulated data set. The average 1-99% interval for the simulated data sets is shown as a shaded box on each plot. (see Figure 4A for a description of kurtosis and skewness measures)
