## Supplemental methods for "A simplified transposon mutagenesis method to perform phenotypic forward genetic screens in cultured cells"

### *Preparation of ligation-mediated PCR templates for Illumina sequencing*

**Enzymatic Fragmentation (300-600 bp Fragments)**

Digest 3 µg of tumor DNA with NEBNext dsDNA Fragmentase enzyme (NEB product #M0348) in a 20 µL volume.

- Notes:
  - Less DNA can be used if concentration is low, but fragmentation times may need to be adjusted.
  - The following protocol differs from supplier's protocol for this enzyme.

1. Combine the following components and mix:

| | |
|---|---|
| 2 µL | 10X Fragmentase Reaction Buffer v2 |
| 1 µL | 200mM $MgCl_2$ |
| x µL | genomic DNA (3 µg) |
| x µL | $H_2O$ |
| **19 µL Total** | |

2. Add 1 µL dsDNA Fragmentase enzyme, vortex to mix, and incubate at room temperature for 10 minutes.
   - Vortex Fragmentase enzyme prior to pipetting

3. Inactivate enzyme by adding 5 µL of 0.5M EDTA

4. Purify the samples using a Qiagen Minelute 96-well clean-up plate (Qiagen product #28053). Resuspend/elute in 25 µL $H_2O$ and shake 5 min. (50 rpm).
   - Alternative DNA clean-up options (e.g. column-based) can be used.

**End Repair**

1. Set up end repair rxn:

| | |
|---|---|
| 25 µL | fragmented genomic DNA |
| 3 µL | 10X NEB CutSmart Buffer (NEB product #B7204) |
| 1 µL | T4 DNA polymerase (NEB product #M0203) |
| 1 µL | 10mM dNTPs |
| **30 µL Total** | |

- Incubate at 12˚C for 15 minutes in thermocycler.

2. Inactivate enzymes by adding 2 µL 160mM EDTA and heating to 75˚C for 20 minutes. Briefly centrifuge plates.

**Ligation**

1. Prepare the adaptors by mixing the linker+ and linker- oligos (each at 100µM in STE Buffer) at a 1:1 ratio (see below for oligo details). Heat to 95˚C for 5 minutes, then allow the oligos to slowly cool to room temperature.

2. Set up ligations:

32 µL fragmented genomic DNA

| | |
|---|---|
| 1 µL | 10X NEB CutSmart Buffer |
| 4 µL | 10 mM ATP |
| 1.5 µL | annealed adaptor |
| 1 µL | T4 DNA ligase (400U; NEB product #M0202) |
| 0.5 µL | 800mM $MgCl_2$ |
| **40 µL Total** | |

Ligate overnight at 16˚C

3. Heat-inactivate the T4 DNA ligase (65˚C for 10 minutes).

4. Digest ligation with *BamHI-HF* overnight at 37˚C. This prevents the fragment from unmobilized transposons from being amplified. *BamHI-HF* solution is made in a 10µL volume per well. To each well add:

   1 µL *Bam*HI-HF
   1 µL NEB CutSmart Buffer
   8 µL $H_2O$
   **10 µL Total**

5. Purify/condense the samples using the Qiagen Minelute 96-well clean-up plate. Resuspend/elute in 25 µL TE and shake 5 min. (50 rpm).
   - Alternative DNA clean-up options (e.g. column-based) can be used.

**Primary PCR:** Two reactions for each sample, one using IRR + Linker primers and one using IRL+ Linker primers.

| | |
|---|---|
| 4.0 µL | ligation reaction |
| 2.5 µL | 10X reaction buffer |
| 1.0 µL | 50mM $MgCl_2$ |
| 0.5 µL | 10 mM dNTPs |
| 0.5 µL | IRR or IRL primer pool (10 µM) |
| 0.5 µL | Linker primer pool (10 µM) |
| 0.1 µL | Platinum Taq polymerase (ThermoFisher product #10966018) |
| 15.9 µL | $H_2O$ |
| 25.0 µL | Total |

| | | |
|---|---|---|
| Step 1 | 94˚C | 2 minutes |
| Step 2 | 94˚C | 30 seconds |
| | 55 to 65˚C | 30 seconds (Increase 1˚C per cycle) |
| | 72˚C | 60 seconds |
| | Repeat 10 cycles | |
| Step 3 | 94˚C | 30 seconds |
| | 65˚C | 30 seconds |
| | 72˚C | 60 seconds |
| | Repeat 25 cycles | |
| Step 4 | 72˚C | 2 minutes |

Hold at 4˚C

SEE NOTE ON PRIMER DESIGNATION FOR PROPER COLOR BALANCING!

- dilute 3 µL of PCR reaction in 72 µL $H_2O$ (1:25 dilution)
- store remaining primary PCR products at 4˚C

**Secondary PCR**

| | |
|---|---|
| 4.0 µL | diluted primary PCR (diluted 1:25 in $H_2O$) |
| 5.0 µL | 10X reaction buffer |
| 1.5 µL | 50mM $MgCl_2$ |
| 1.0 µL | 10 mM dNTPs |
| 1.0 µL | Nextera Illumina Primers (Non-dilute, straight from plate; 13.3uM) |
| 0.2 µL | Platinum Taq polymerase |
| 37.3 µL | $H_2O$ |
| 50.0 µL | Total |

- perform PCR using the same cycling conditions as primary PCR, but with 20 cycles instead of 25 for step 3

- Analyze 15 µL of PCR product on 1.5% agarose gel. Each sample should appear as a smear of DNA with maximum intensity between ~150-500bp. If specific bands are detected for a sample, try repeating the primary and/or secondary PCR.
- Purify remaining PCR product to remove excess primers/dNTPs (e.g. with Qiagen's Minelute 96 UF PCR purification kit).
- Determine concentration of purified PCR products (e.g. with Nanodrop)
- Pipet 200 ng of each PCR product pool into a single tube to be run on a single lane on the Illumina platform
- Adjust the final concentration of the mixed sample to be ~20-25 ng/µl (May vary based on sequencer used for run)
- Incubate the diluted products at 37-42˚C for 30 minutes
- Submit sample for sequencing

**Oligos to generate adaptors:**

Linker+ `5'-GTAATACGACTCACTATAGGGCTCCGCTTAAGGGAC-3'`
Linker- `5'-Phos-GTCCCTTAAGCGGAG-C3spacer-3'`

Adaptor oligos are resuspended in STE* buffer at 100µM. All PCR primers were used at 10 µM concentration. C3spacer modification is available from Integrated DNA Technologies.

**Primers for IRR amplification (recognition bases only):**

<u>Primary PCR</u>
IRR primer `5' GGATTAAATGTCAGGAATTGTGAAAA 3'`
Linker primer `5' GTAATACGACTCACTATAGGGC 3'`

**Primers for IRL amplification (recognition bases only):**

Primary PCR

IRL primer 5' AAATTTGTGGAGTAGTTGAAAAACGA 3'

Linker primer 5' GTAATACGACTCACTATAGGGC 3'

Additionally, each primary PCR primer has an Illumina tag on the 5' end that recognizes the secondary PCR Illumina primers (Blue). Between the Illumina tag and recognition bases (shown above, Red) there are stuffer bases for color balance (Black). Below is a representative primer.

IRL_V1.1:

TCGTCGGCAGCGTCAGATGTGTATAAGAGACAGHAAATTTGTGGAGTAGTTGAAAAACGA

Each primer (IRL, IRR, and linker) has five different stuffer base sequences (denoted H above). These five primers are mixed together at an equal concentration (20ul of each for a total volume of 100ul at 100uM concentration stock, designation IRL_V1 Pool for example).

Here is a list of all the primary primers:

IRL_V1 POOL

*IRL_Nextera_V1.1*

*TCGTCGGCAGCGTCAGATGTGTATAAGAGACAGHAAATTTGTGGAGTAGTTGAAAAACGA*

*IRL_Nextera_V1.2*

*TCGTCGGCAGCGTCAGATGTGTATAAGAGACAGNCAAATTTGTGGAGTAGTTGAAAAACGA*

*IRL_Nextera_V1.3*

*TCGTCGGCAGCGTCAGATGTGTATAAGAGACAGNNYAAATTTGTGGAGTAGTTGAAAAACGA*

*IRL_Nextera_V1.4*

*TCGTCGGCAGCGTCAGATGTGTATAAGAGACAGNNYCAAATTTGTGGAGTAGTTGAAAAACGA*

*IRL_Nextera_V1.5*

*TCGTCGGCAGCGTCAGATGTGTATAAGAGACAGNNNBCCAAATTTGTGGAGTAGTTGAAAAACGA*

IRR_V1 POOL

*IRR_Nextera_V1.1*

*TCGTCGGCAGCGTCAGATGTGTATAAGAGACAGHGGATTAAATGTCAGGAATTGTGAAAA*

*IRR_Nextera_V1.2*

*TCGTCGGCAGCGTCAGATGTGTATAAGAGACAGNNGGATTAAATGTCAGGAATTGTGAAAA*

*IRR_Nextera_V1.3*

*TCGTCGGCAGCGTCAGATGTGTATAAGAGACAGNNYGGATTAAATGTCAGGAATTGTGAAAA*

*IRR_Nextera_V1.4*

*TCGTCGGCAGCGTCAGATGTGTATAAGAGACAGNNNYGGATTAAATGTCAGGAATTGTGAAAA*

*IRR_Nextera_V1.5*

*TCGTCGGCAGCGTCAGATGTGTATAAGAGACAGNNNYCGGATTAAATGTCAGGAATTGTGAAAA*

LINK_V1 POOL

link_Nextera_V1.1

TCGTCGGCAGCGTCAGATGTGTATAAGAGACAGGTAATACGACTCACTATAGGGC

link_Nextera_V1.2

TCGTCGGCAGCGTCAGATGTGTATAAGAGACAGHNGTAATACGACTCACTATAGGGC

link_Nextera_V1.3

TCGTCGGCAGCGTCAGATGTGTATAAGAGACAGNNYBGTAATACGACTCACTATAGGGC

link_Nextera_V1.4

TCGTCGGCAGCGTCAGATGTGTATAAGAGACAGNNYBCNGTAATACGACTCACTATAGGGC

link_Nextera_V1.5

TCGTCGGCAGCGTCAGATGTGTATAAGAGACAGNNYBCNCCCGTAATACGACTCACTATAGGGC

**Secondary PCR (for IRR and IRL)**
We are using the Nextera XT index primers.  Here are the product #'s:
FC-131-2001
FC-131-2002
FC-131-2003
FC-131-2004
We ordered the primers from IDT at a discounted rate. Your DNA sequencing facility may also have Illumina index primers on hand for investigator use.

*STE Buffer
50 mM NaCl
10 mM Tris-Cl (pH 8.0)
1mM EDTA (pH 8.0)
